# A two-trick pony: lysosomal protease cathepsin B possesses surprising ligase activity

**DOI:** 10.1101/2020.05.07.082842

**Authors:** Tyler R. Lambeth, Zhefu Dai, Yong Zhang, Ryan R. Julian

**Affiliations:** Department of Chemistry, University of California, Riverside, California 92521, United States; Department of Pharmacology and Pharmaceutical Sciences, School of Pharmacy, Department of Chemistry, Dornsife College of Letters, Arts and Sciences, Norris Comprehensive Cancer Center, and Research Center for Liver Diseases, University of Southern California, Los Angeles, California 90089, United States

**Keywords:** cathepsin B (catB), protease, ligase, lysosome, occluding loop

## Abstract

Cathepsin B is an important protease within the lysosome, where it helps recycle proteins to maintain proteostasis. It is also known to degrade proteins elsewhere but has no other known functionality. However, by carefully monitoring peptide digestion with liquid chromatography and mass spectrometry, we observed synthesis of novel peptides during cathepsin B incubations. This ligation activity was explored further with a variety of peptide substrates to establish mechanistic details and was found to operate through a two-step mechanism with proteolysis and ligation occurring separately. Further explorations using varied sequences indicated increased affinity for some substrates, though all were found to ligate to some extent. Finally, experiments with a proteolytically inactive form of the enzyme yielded no ligation, indicating that the ligation reaction occurs in the same active site but in the reverse direction of proteolysis. These results clearly establish that cathepsin B can act as both a protease and ligase, although protease action eventually dominates over longer periods of time.

## Introduction

Cathepsin B (catB) is a cysteine protease and one of several members of the cathepsin family, which are localized primarily in lysosomes. The cathepsins serve to digest proteins and peptides sent to the lysosome for recycling, breaking down substrates into their constituent amino acids which are released for future synthesis or energy production. Like most of the cathepsin family, catB possesses endoprotease activity, allowing it to cleave between internal residues in a protein sequence. However, catB is best known for its exopeptidase activity, where it preferentially removes two amino acids at a time from the C-terminus of a peptide substrate. This uncommon dual functionality among the cathepsins is due to the presence of an occluding loop comprised of 20 amino acids which blocks one end of the active site and restricts the space to accommodate only two residues past the catalytic cysteine.^1^ This unique two-residue slot is shown adjacent to the active site and catalytic residue of catB in Fig. 1.^2,3^ In addition to steric allowances, the occluding loop contains two histidine residues at positions 110 and 111 that bind to the properly positioned C-terminal carboxylate and facilitate exopeptidase activity.

**Figure 1.**
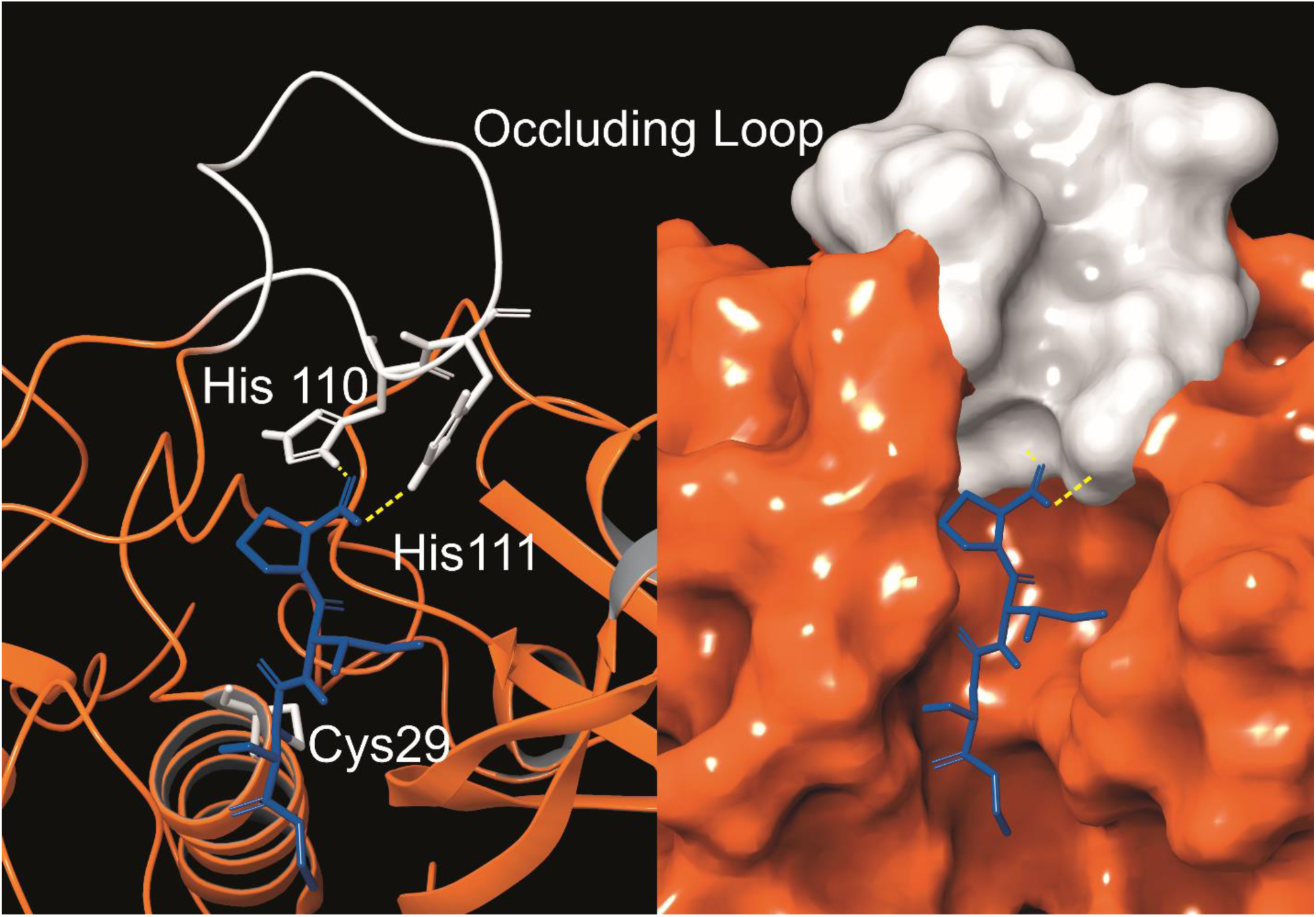
The active site of catB shown with a bound substrate inhibitor in ribbon representation (left) and molecular surface representation (right). The occluding loop and catalytic cysteine are shown in white, and the substrate is shown in blue. His110 and His111 from the occluding loop bind to the C-terminus of a substrate to favor carboxypeptidase activity. Images constructed from PDB ID:1CSB.

In addition to the lysosome, catB is found in a variety of cellular locations where it is involved in proteolysis. One secondary role is the activation of enzymes such as trypsin, ß-Galactosidase, and renin after synthesis of the respective proenzymes.^4^ In antigen presenting cells, catB is thought to be involved in pruning sequences along with other cathepsins in endosomes before selection by the major histocompatibility complex.^5^ CatB has also been implicated in apoptosis, where its release into the cytosol after lysosomal membrane permeabilization cleaves the apoptotic protein Bid into its active form.^6^ Another role catB plays outside the lysosome is degradation of the extracellular matrix, often to promote tumor growth in cancer.^7^ Like other lysosomal proteases, catB is most active at acidic pH, however it can function at near-neutral pH in the cytosol.^8^ Due to the loss of salt-bridges required for stabilization of the occluding loop, catB operates primarily as an endopeptidase at higher pH. These secondary roles for catB demonstrate its versatility but do not reveal any known function outside proteolysis.

While peptide bonds are facile to cleave using proteases or chemical methods such as acid hydrolysis^9^, synthesis of new bonds in aqueous solution is a more difficult challenge. Due to their high stability, carboxylic acids are not naturally reactive enough for amines to attack and spontaneously form a peptide bond. Instead targets must be modified into a more reactive form first. Biologically, new peptides are formed in the ribosome using an active ester as a target for a nucleophilic amine attack, binding the N-terminal amine from an amino acid to the carbonyl from the C-terminus of the growing peptide chain.^10^ In vitro methods commonly exploit this same type of ester-amine reaction, most famously in the solid-phase peptide synthesis and NHS-ester reactions.^11^ Ester-amine reactions are non-specific to the termini however, and unlike ribosomal synthesis which controls the reaction sterically, any free amine such as a lysine can form an incorrect linkage. For this reason, these reactions require careful protection and subsequent de-protection of the side-chains to increase the reaction efficiency. Native chemical ligation is an alternative to standard ester-based reactions, utilizing a thioester and a cysteine instead.^12^ It has the advantage of operating orthogonally to canonical amino acids, so no protection of peptide side-chains is required. However, this method still requires chemical modification to produce a reactive C-terminal thioester group which is not found naturally in peptides, and the reaction is also limited to sites with a cysteine residue. In biology, ligation of existing peptides is performed by enzymes known as ligases, which catalyze connections between the N-terminus and C-terminus.^13^ These ligases are rare however, and typically highly specific in their use, often for antigen preparation for the immune system.^14^ An ideal replacement for these options would maintain the high specificity of enzymes while eliminating side-products or the need for protecting or reactive groups.

Herein, we examine incubations of catB with synthetic peptide sequences via liquid chromatography and mass spectrometry (LC-MS) to evaluate its ability to perform previously unreported ligation reactions between peptide substrates. Ligated products were detected in multiple incubations, and we explored the sequence specificity for the reaction by substituting amino acids with different functional groups. Additional experiments using a mutated and non-proteolytic form of the enzyme revealed the role of the active site residues and provide further information about the ligation mechanism.

### Experimental Procedures

#### Peptide Synthesis

All peptides were synthesized following an accelerated FMOC-protected solid-phase peptide synthesis protocol.^11^ Following synthesis, peptides were purified on a Varian ProStar HPLC using a Phenomenex Jupiter Proteo C12 4 μm 90 Å 250 mm × 4.6 mm column and then validated by LC-MS. Purified peptides were stored frozen in water and vacuum-dried before reconstitution in appropriate buffers.

#### Cathepsin B Incubations

Cathepsin B for initial incubations was purchased from Athens Research. Purified peptides were dissolved in 50 mM acetate buffer pH 5.5 to a final concentration of 0.5 mg/mL with 1 mM EDTA and 1 mM DTT to prevent active site oxidation. Timepoint aliquots were removed after 2 hours and diluted to a final concentration of 5 µM with 0.2% TFA solution to pH quench the enzymatic reaction. Digested samples were frozen until analysis.

#### LC-MS Analysis

Samples were analyzed on an Agilent 1100 HPLC interfaced with a ThermoFisher LTQ using an electrospray ionization source. During HPLC, peptides were separated on a ThermoScientific BetaBasic C-18 3 µM, 150 x 2.1 mm column using water with 0.1% formic acid for mobile phase A, and acetonitrile with 0.1% formic acid for mobile phase B. Samples were separated with a gradient of 5-70% phase B over 32.5 minutes.

## Results and Discussion

While evaluating LC-MS results for the stepwise proteolysis of a synthetic peptide APSWFDTGLSEMR, an unusual product corresponding to excision of an internal sequence (APSWFDTGMR) was observed as shown in Fig. 2a. This result appeared to suggest that the lysosomal protease catB had removed the internal residues LSE from the peptide or had otherwise cleaved the peptide multiple times and stitched the ends back together. Careful evaluation confirmed that the APSWFDTGMR product was not present in the initial sample and must have been produced during incubation, as shown by the data in Fig. S1. The identity of this product was further examined by collision-induced dissociation (CID), yielding characteristic b and y ions covering most of the sequence as shown in Fig. 2b. Importantly, the b8 and b9 ions map out the unexpected C-terminal linkage, definitively confirming the proposed ligation product. Generation of this new ligated peptide product requires initial cleavage of peptide bonds 8 and 11, followed by creation of a new peptide bond between Gly8 and the methionine in MR. The act of rejoining peptide fragments, or apparent peptide ligase activity, has not been previously reported for catB. Surprisingly, the ligation product comprises almost 20% of the detected peaks after 2 hours of incubation, indicating that ligation can be reasonably favorable under appropriate conditions.

**Figure 2.**
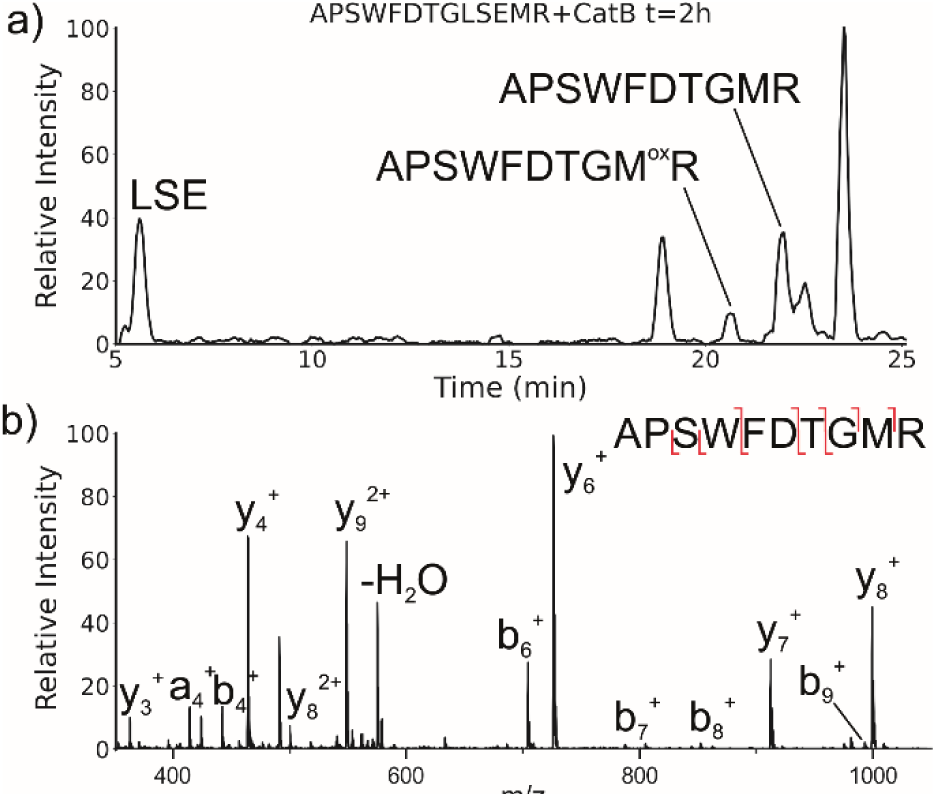
(a) LC chromatogram for the incubation of APSWFDTGLSEMR with catB. The labelled fragment APSWFDTGMR is produced by protease cleavage first followed by ligation of the dipeptide MR onto the proteolytic fragment APSWFDTG. (b) CID fragmentation of the ligated peak APSWFDTGMR.

While this behavior has previously gone unnoticed in catB, dual protease/ligase ability has been observed in the enzyme legumain.^15^ Legumain participates in the MHC antigen cleaving pathway but can also ligate cyclic peptides that are involved in plant defense mechanisms. Legumain ligation operates through a reactive succinimide not directly associated with the proteolytic cysteine. Instead, the succinimide lies directly adjacent to the histidine of the catalytic triad, which allows it to share the same active site as the proteolytic machinery. For catB, ligase activity appears to target dipeptides such as MR, which may suggest involvement of the same occluding loop and catalytic site responsible for exopeptidase action. Ligase behavior would simply constitute reversal of the protease action as illustrated in Scheme 1.

**Scheme 1.**
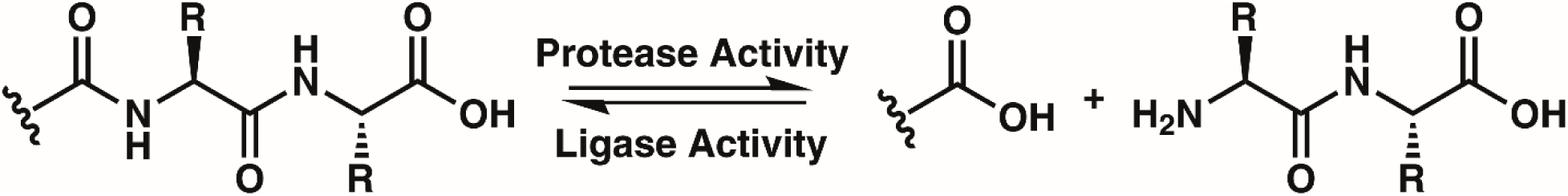
The protease activity of CatB removes two amino acid residues as a dipeptide from the C-terminus of a substrate, while the ligase activity attaches a dipeptide to the C-terminus of a substrate.

To elucidate whether the internal removal and ligase activity occurs in one step or multiple steps, two synthetic sequences, APSWFDTGLSEMR and GPSWFDTGLSEGR were incubated with catB as shown in Fig. 3a. While a one-step process would yield the original sequences minus the internal fragment, a two-step process could lead to scrambling of the C-terminal appendages. The color-coded sequences shown in Fig. 3 confirm that scrambling occurred, suggesting that the two C-terminal amino acids are first removed and released, followed by subsequent reattachment following removal of LSE. This illustrates not only that the process occurs in two steps, but also that it is not sequence-specific to the MR portion of the original APSWFDTGLSEMR peptide. Quantitation of the homogenous ligated products which are comprised of only the original sequence versus the heterogeneous products comprised of mixed sequences is shown in Fig. 3b. The amount of homogenous ligation is significantly higher, perhaps due to the nearby localization of both necessary pieces after proteolysis.

**Figure 3.**
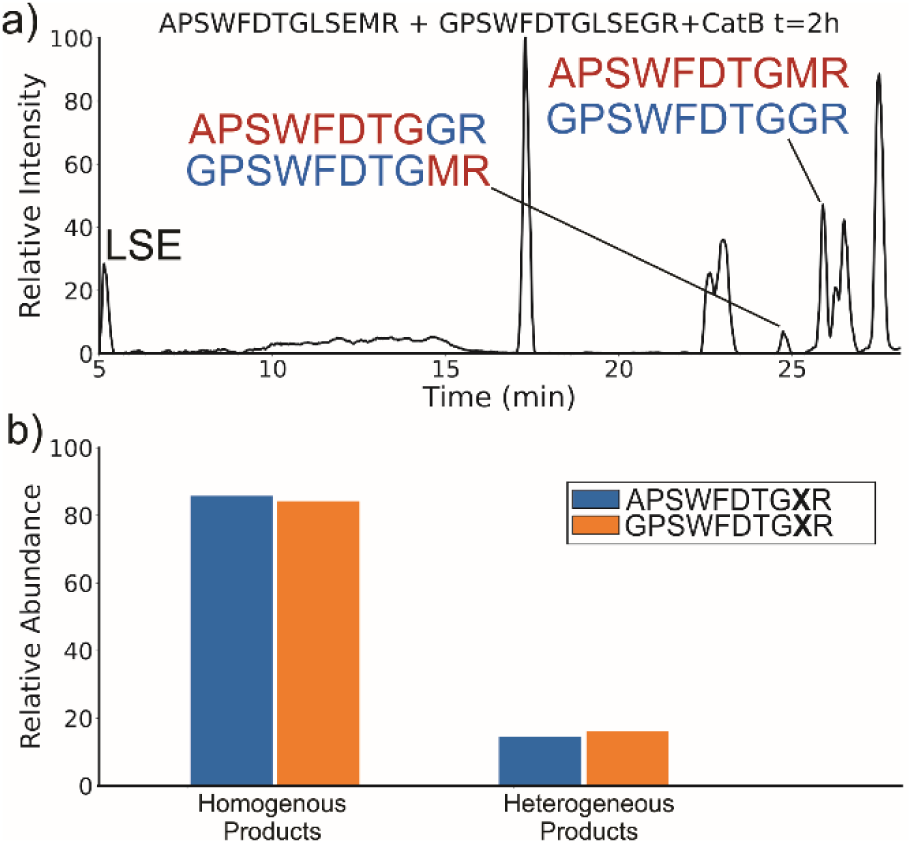
(a) LC chromatogram of the incubation of APSWFDTGLSEMR and GPSWFDTGLSEGR with CatB. Multi-colored labels indicate products formed by ligation of fragments from different starting sequences. (b) Relative quantitation of the homogenous versus heterogeneous ligation products.

To further explore the sequence specificity, modifications were made to the C-terminal residue to determine if it operates as a key factor in the ligase activity (see Fig. 4, derived from data in Fig. S2). The sequences APSWFDTGLSEMR and GPSWFDTGLSEGR were incubated with the synthetic dipeptides GK and GA, to evaluate whether arginine or any basic residue is required. In this incubation, ligation was detected for both the lysine and alanine substitutions, indicating that neither arginine nor a basic residue is a necessary element in ligation. With this knowledge, an incubation of both dipeptides with a new sequence, VFFAEDVGSNK, was performed and is shown in Fig. 5. In this incubation, the full sequence with appended dipeptide GA was detected, as well as the proteolytic fragment VFF with ligated dipeptides GA and GK in separate peaks. These fragments, especially the appended full sequence, demonstrate that the previous internal remove of LSE before ligation is a coincidental action resulting from competing protease activity, and that proteolysis is not required for ligase activity. The low abundance of these fragments also shows that while the ligase activity ostensibly works for a variety of sequences, it appears to be affected by substrate affinity. This illustrates why ligase activity may have gone unnoticed previously, as the initial incubation substrate APSWFDTG appears to have a high affinity for ligase activity, allowing for easy detection while other ligation products are harder to detect. The competing protease activity also serves to break down any ligated products, and by the end of typical incubations lasting 12 hours or more, these sequences have been fully digested, erasing the evidence. No fragments larger than two residues were attached to the C-terminus of a peptide in any incubation, further implying that the occluding loop may play a role in ligation. Experiments with isomeric substitutions were conducted to further probe the limits of catB ligation. An incubation of catB with the L-isoAsp isomer of APSWF<u>D</u>TGLSEMR resulted in the ligated fragment FDTGMR, as shown in Fig. 6a. The identity of the fragment was confirmed with CID fragmentation as shown in Fig. S3. This ligation product is also obtained by removal of the internal LSE sequence, however this product does not appear in the incubation with the canonical sequence shown in Fig. 2, which does not show any abundant cleavage at the W-F bond. W-F cleavage results seemingly from the recognition of the isoAsp side-chain as a C-terminus. The altered connectivity of L-isoAsp reduces the side-chain length by one carbon while similarly lengthening the backbone connection. This new orientation produces a structural analogue of a normal C-terminal carboxylate group, as illustrated in Fig. 6b. When this C-terminal mimic is bound by catB, the F<u>D</u> residues from APSWF<u>D</u>TGLSEMR sit in the pocket normally reserved for the last two residues of a sequence, and cleavage occurs at the W-F bond as it would to remove a dipeptide. This unintended behavior has not been previously reported for catB but has been noted for other carboxypeptidases.^16^

**Figure 4.**
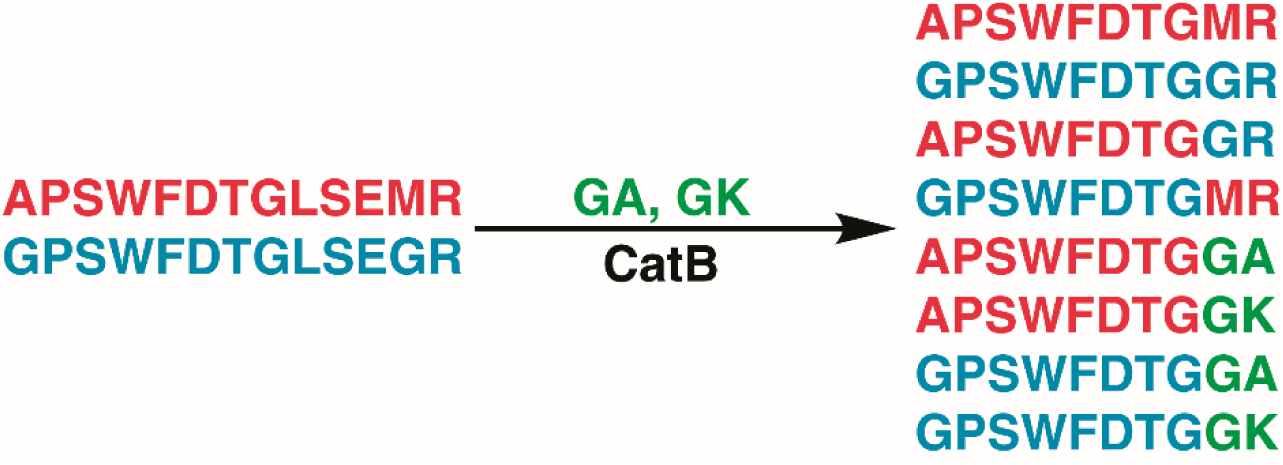
Incubation of APSWFDTGLSEMR and GPSWFDTGLSEGR with dipeptides GA and GK results in a series of heterogeneous and homogenous ligation products. Raw LC chromatogram shown in Supporting Information Figure S1.

**Figure 5.**
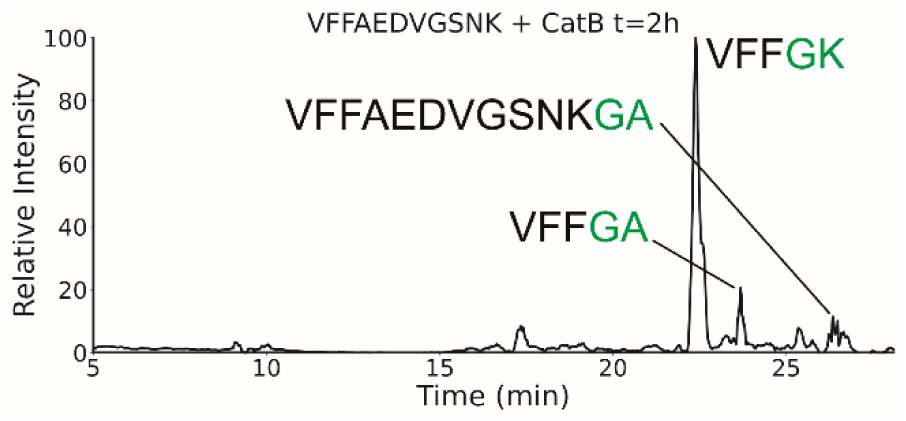
Extracted ion chromatogram for the incubation of VFFAEDVGSNK with CatB. Ligation is detected on proteolytic fragments as well as the full intact sequence.

**Figure 6.**
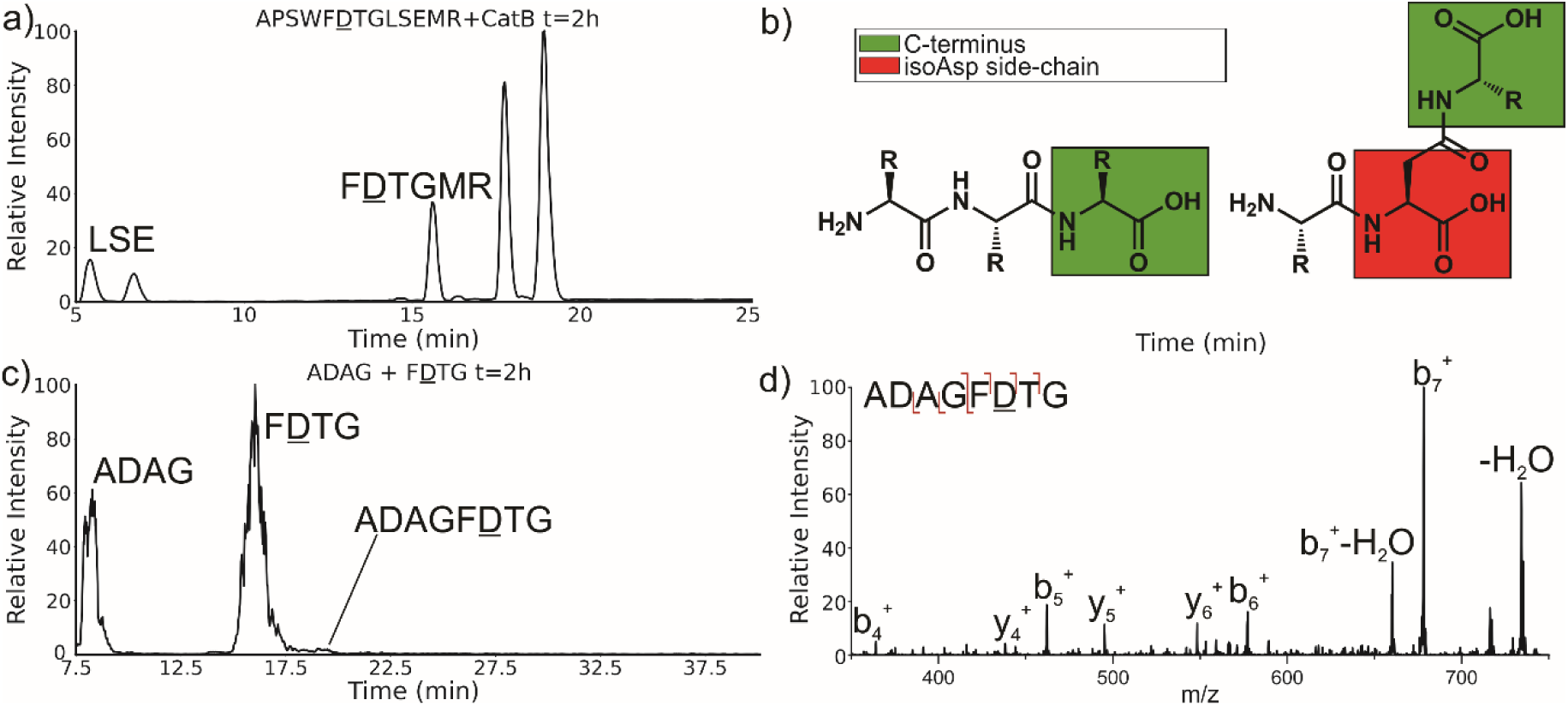
(a) LC chromatogram of the incubation of APSWF<u>D</u>TGLSEMR. Incubation produces the ligated fragment F<u>D</u>TGMR which has been cleaved near the L-isoAsp residue due to recognition by catB. (b) Structures of a canonical peptide (left) and L-isoAsp containing peptide (right). Due to the altered connectivity of L-isoAsp, the side-chain structurally resembles a C-terminus and can be recognized by catB as a binding target. (c) LC chromatogram for the incubation of ADAG and F<u>D</u>TG with catB. The incubation produces the ligated fragment ADAGF(isoAsp)TG, CID fragmentation shown in (d).

The ability of catB to recognize L-isoAsp as a C-terminal residue suggests that it could also serve as a site for ligation. In order to explore this possibility, two sequences were incubated together with catB, ADAG and F<u>D</u>TG, modelled after the ligated fragment from Fig. 6a. Indeed, after a period of incubation followed by LC-MS analysis, the ligated fragment ADAGF<u>D</u>TG was identified in a small quantity as shown in Fig. 6c. This peptide is the first ligation product that does not correspond to attachment of a simple dipeptide. In this case, the first two residues of F<u>D</u>TG take on the role of a dipeptide and are ligated together with ADAG. These results reveal that a peptide with an L-isoAsp in the second position can be attached to another peptide regardless of the size limitation normally imposed by the occluding loop. As L-isoAsp can be easily repaired to L-Asp by the repair enzyme Protein L-isoaspartyl methyltransferase,^17^ this method of utilizing L-isoAsp to ligate longer sequences could be used to produce longer canonical peptide sequences.

The implied participation of the occluding loop by preferential ligation of dipeptides (or dipeptide mimics) suggests that the same active site may be used for both proteolysis and ligation in catB. To probe whether the same active site is required for ligation, incubations were performed with a mutated form of catB. The C29A mutant lacks the catalytic cysteine required for proteolysis, as demonstrated previously.^18^ Incubation of catB-C29A with APSWFDTG and the dipeptide MR is shown in Fig. 7a. This incubation did not yield any ligated APSWFDTGMR products when investigated by LC-MS analysis. Within the same timeframe, wild type catB was able to produce the necessary fragments (APSWFDTG and MR) via proteolysis of APSWFDTGLSEMR, and then ligate them together again into the target peptide APSWFDTGMR shown in Fig. 7b. The results in Fig. 7b also represent a replicate of those shown in Fig. 2a, with the exception of a completely different source of catB. Overall, the results indicate that the proteolytic cysteine is necessary for ligation and provide evidence that ligation operates by reversal of the proteolytic mechanism.

**Figure 7.**
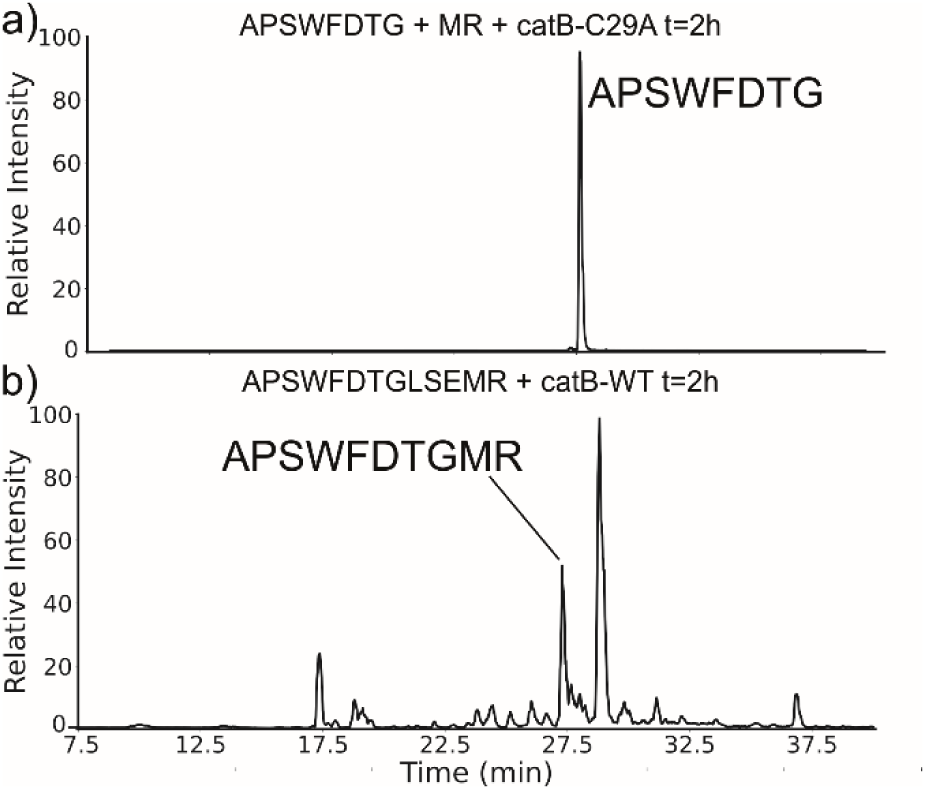
(a) LC chromatogram of the incubation of APSWFDTG and peptide MR with catB-C29A mutant. No amount of ligated product was detected. (b) Incubation of the peptide APSWFDTGMR with wild type catB produces the ligated peak APSWFDTGMR in the same timeframe.

## Conclusions

In summary, unique ligase activity was identified in catB, which was previously only recognized as a protease. Explorations of this reaction revealed that it has broad sequence tolerance though varied substrate affinity. Dipeptides are preferentially coupled to the C-terminus in the ligation, but we demonstrate that catB can produce sequences of any length by utilizing an isoAsp residue as a C-terminal mimic. A highly specific ligation reaction such as this could be used to produce specific target sequences, to couple protein domains together, or to add modifications such as fluorescent tags to existing peptides without any supporting chemical reactions or protecting groups. Future experiments could optimize the ligase function of catB by mutagenesis of the residues in close proximity to the active-site. More extensive substrate libraries could also be screened with catB to determine relative propensities for ligation, opening up new possibilities for targeted substrate design for in vitro and in vivo ligations.

## Funding and additional information

The authors are grateful for funding from the NIH (1R01AG066626-01 for RRJ and 1R35GM137901-01 for YZ).

