## additional data for "A two-trick pony: lysosomal protease cathepsin B possesses surprising ligase activity"

### Supporting Information

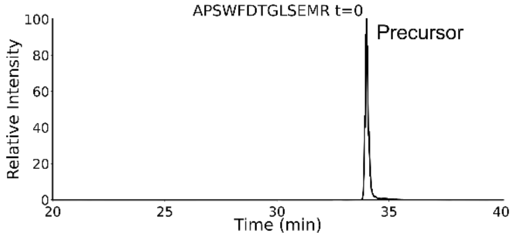


**Figure S1.** LC chromatogram for the initial time point of APSWFDTGLSEMR + CatB. No peaks were present except for the precursor peptide.

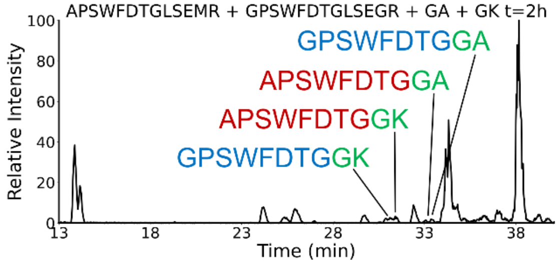


**Figure S2.** LC chromatogram for the incubation of APSWFDTGLSEMR and GPSWFDTGLSEGR with dipeptides GA and GK with CatB. Raw data for the results shown in Scheme 2.

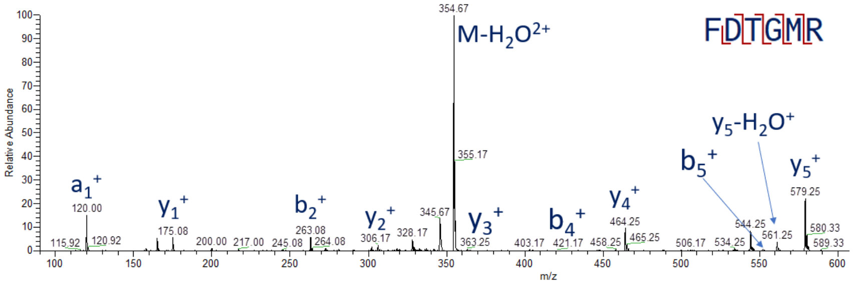


**Figure S3.** CID fragmentation spectrum for the ligation product FDTGMR.
